# Scalable and accurate automated method for neuronal ensemble detection in spiking neural networks

**DOI:** 10.1101/2020.10.12.335901

**Authors:** Rubén Herzog, Arturo Morales, Soraya Mora, Joaquin Araya, María-José Escobar, Adrian G. Palacios, Rodrigo Cofré

## Abstract

We propose a novel, scalable, and accurate automated method for detecting neuronal ensembles from a population of spiking neurons. Our approach offers a simple yet powerful tool to study ensemble activity. It allows the participation of neurons in different ensembles, has few parameters to tune and is computationally efficient. We used spike trains of retinal ganglion cells obtained from multi-electrode array recordings under a simple ON-OFF light stimulus to test our method. We found a consistent stimuli-evoked ensemble activity intermingled with spontaneously active ensembles and irregular activity. Our results suggest that the early visual system activity is already organized in clearly distinguishable functional ensembles. To validate the performance and generality of our method, we generated synthetic data, where we found that our method accurately detects neuronal ensembles for a wide range of simulation parameters. Additionally, we found that our method outperforms current alternative methodologies. Finally, we provide a Graphic User Interface, which aims to facilitate our method’s use by the scientific community.

**Author summary:** Neuronal ensembles are strongly interconnected groups of neurons that tend to fire together (Hebb 1949). However, even when this concept was proposed more than 70 years ago, only recent advances in multi-electrode arrays and calcium imaging, statistical methods, and computing power have made it possible to record and analyze multiple neurons’ activities spiking simultaneously, providing a unique opportunity to study how groups of neurons form ensembles spontaneously and under different stimuli scenarios. Using our method, we found that retinal ganglion cells show a consistent stimuli-evoked ensemble activity, and, when validated with synthetic data, the method shows good performance by detecting the number of ensembles, the activation times, and the core-cells for a wide range of firing rates and number of ensembles accurately.

## 1 Introduction

Donald Hebb predicted more than 70 years ago that ensembles would naturally arise from synaptic learning rules, where neurons that fire together would wire together [1]. However, despite the long history of this idea, only recently the simultaneous recordings and computational analysis from hundreds of cells have turned out to be possible [2]. Recent advances in recording technology of neuronal activity combined with sophisticated methods of data analysis have revealed significant synchronous activity between neurons at several spatial and temporal scales [3–5]. Theses groups of neurons that have the tendency to fire together known as *neuronal ensembles* (also called cell assemblies) are hypothesized to be a fundamental unit of neural processes and form the basis of coherent collective brain dynamics [1, 6–8].

The idea that the coding units of information are groups of neurons firing together (not single neurons) represented a paradigm shift in the field of computational neuroscience [6]. Neuronal ensembles have been proposed as a fundamental building block of the whole-brain dynamics, and relevant to cognitive functions, in particular, as ensemble activity could implement brain-wide functional integration and segregation [3].

Large-scale neuronal recordings techniques such as multi-electrode arrays (MEA) or calcium imaging, allow for the recording of the activity of hundreds and even thousands of neurons simultaneously [5, 9–12]. These recent technological advances provide a fertile ground for analyzing neuronal ensembles and investigating how collective neuronal activity is generated in the brain. Recent studies using multi-neuronal recording techniques have revealed that a hallmark of population activity is the organization of neurons into ensembles, generating new insights and ideas about the neural code [9, 13–15]. In particular, the activation of specific ensembles has been shown to correlate with spontaneous and stimuli evoked brain function [16]. The brain-wide alterations present in neurological and mental impairments disrupt population activity patterns and therefore affect the neuronal ensembles. Indeed, neuronal ensembles are susceptible to epileptic seizures and schizophrenia as shown in *in vivo* two-photon calcium imaging data in mouse [17, 18], in medically-induced loss of consciousness in mice and human subjects [19] and in a mice model of autism [20].

However, identifying and extracting features of ensembles from high-dimensional spiking data is challenging. Neuronal ensembles have different sizes and have different activity rates. Some neurons may not participate in ensemble activity, while others may participate in many, and not all neurons within an ensemble fire when the ensemble is active. Ensembles can exhibit temporal extension, overlap, or display a hierarchical organization, making it difficult to distinguish between them [2].

Sophisticated statistical and computational tools are needed to extract relevant features of neuronal activity. Detecting neuronal ensembles can be posed as a clustering problem, where the binary population patterns (one-time bin of the spike train) generated by the neural population are the variables to cluster. Given the probabilistic nature of spiking activity, the challenge is to distinguish between core and non-core cells, i.e., cells that belong to the ensemble and cells that do not, respectively.

There exists a wide variety of techniques and ideas that have been used to detect and interpret neuronal ensembles (see Refs. [13, 21, 22] for reviews), for example, previous works have applied different methodologies such as principal component analysis [23–26], correlation between neurons [26–28], correlation between population spiking patterns [16], statistical evaluation of patterns [29–32] and non-negative matrix factorization [25, 33]. Among them, only the one based on the correlation between population patterns, and the one based on non-negative matrix factorization, fit with a definition of ensembles. Under this definition, neurons may participate in many ensembles, and ensembles have a specific duration in time. However, these methods are computationally expensive, and require tuning several parameters, which hinder their application by the scientific community.

We exemplify our method’s advantages on spiking neuronal data recorded using MEA from mouse *in vitro* retinal patch, from which the spiking activity of hundreds of retinal ganglion cells (RGCs) is obtained during a simple ON-OFF stimuli, i.e. consequtive changes between light and darkness.

In brief, the vertebrate retina is part of the central nervous system composed of thousand of neurons of several types [34–36], organized in a stratified way with nuclear and plexiform layers [37]. This neural network has the capability to process several features of the visual scene [38], whose result is conveyed to the brain through the optic nerve, a neural tract composed mainly by the axons of the RGCs. In fact, the physiological mechanisms involved in many of those processes are starting to become clear with the development of new experimental and computational methods [36, 37, 39]. One of the most remarkable, and simple, example of retinal processing is the ON-OFF responses of RGCs, a stereotypical increase or decrease in firing rate when confronted to changes light intensity. In this case, the connectivity between RGCs and Bipolar cells (Bc) in the Inner Plexiform Layer (IPL) plays a major role, determining the tendency of RGC to preferentially fire when the light increased, decreased, or both [37]; this property is often called polarity [40], and represents the broadest functional classification of RGCs into ON, OFF, and ON-OFF cell types.

We hypothesize that retinal ensembles could also exhibit this property as a whole. Our analysis revealed the existence of diverse ON and OFF retinal ensembles with a specific stimulus preference as functional units, which suggests that a stimulus tuning preference is a property of the ensembles as a whole, and not a simple inheritance from their corresponding core-cells. Besides, we validated our method on synthetic data where ensembles were artificially generated, showing a remarkable detection performance for the ensemble number, the ensemble activation sequence, and the core-cells detection over a wide range of parameters.

Thus, we tested and validated our method using biological and synthetic data, showing its accuracy and broad applicability to different scenarios. To facilitate our method’s use by the community, we provide a Graphic User Interface and the codes that implement our algorithm that aim to provide a fast, scalable, and accurate solution to the problem of detecting neuronal ensembles in multi-unit recordings.

## 2 Results

### Detection of ensembles on RGC under a simple ON-OFF stimulus

We show our method’s usefulness on a parallel recording of a mouse retinal patch *in vitro* using MEA. We analyzed the spike response of retinal ganglion cells (RGCs) under a simple ON-OFF light stimulus, where neuronal ensembles are detected without prior information about the stimulus. Their functional role is evaluated in terms of stimulus tuning preference. As expected for RGCs, the sum of all the emitted spikes in a given time bin (population activity) is tightly locked to the stimulus, transiently increasing each time the stimulus changes (Fig. 1A, top panel). After this harsh response, the population activity decays exponentially until it reaches a stable point. However, the population activity evoked by the ON-stimulus is different in amplitude and shape from the ones evoked by the OFF-stimulus.

**Fig 1.**
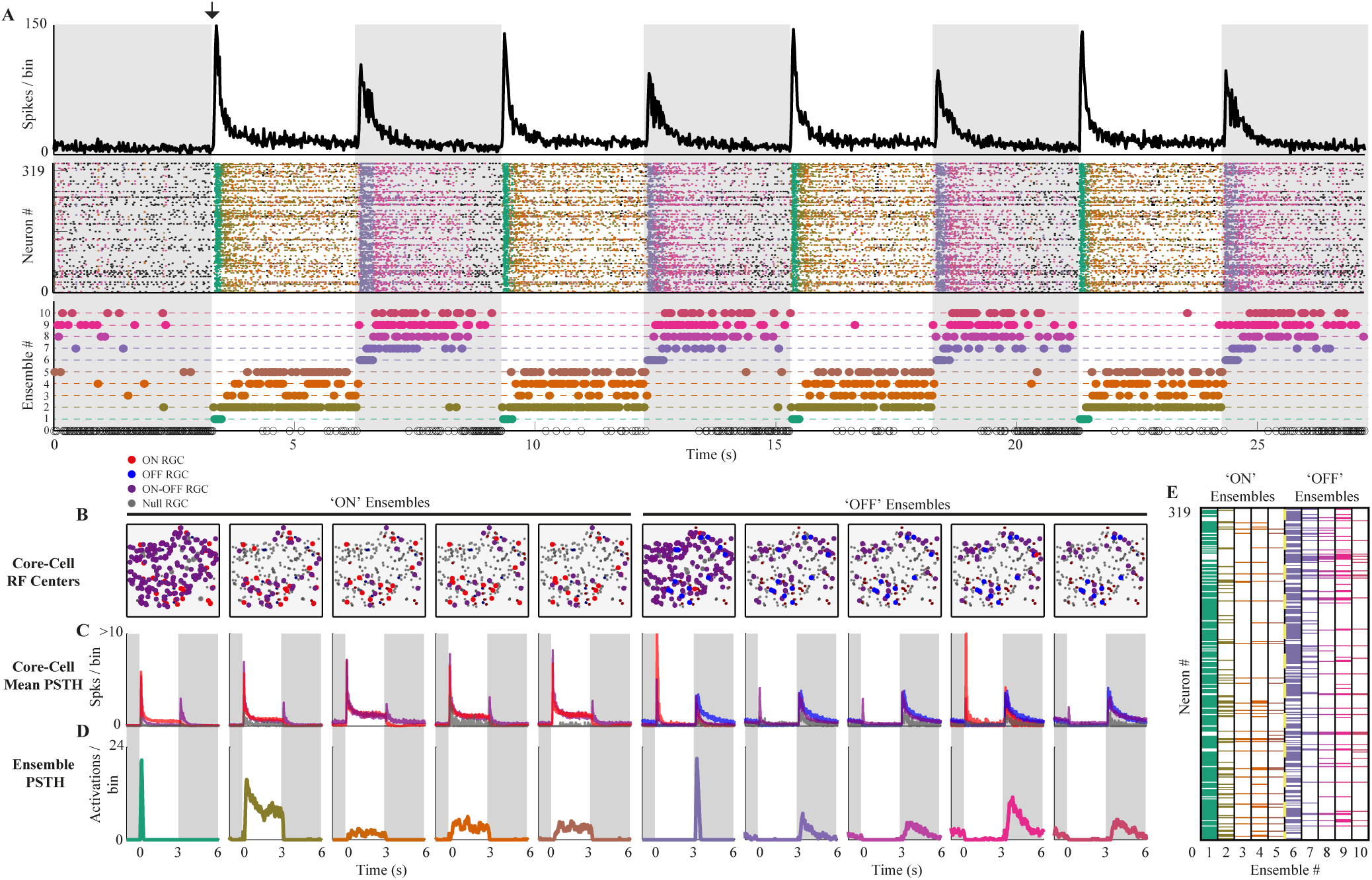
Stimulus-evoked retinal ensembles. **(A) Top.** Population activity (spikes per bin) of 319 RGC’s in time during ON light stimulation (white background) and OFF (gray background). The black arrow shows the ON stimulus started. As expected for the retina, the population activity transiently increases when the stimulus switches and then decays and stabilizes. **Middle.** Each spiking pattern is colored according to the ensemble to which they belong. **Bottom.** The activation of each ensemble in time is represented by a colored dot (matching the spike pattern color code). The method detects a consistent stimulus-locked ensemble activity with ON (ensembles from 1 to 5) and OFF (ensembles from 6 to 10) ensembles. Ensemble 0 represents all the spiking patterns that were discarded according to rejection criteria (see Methods). **(B)** The estimated receptive field centers of all the RGCs are shown. The color indicates the RGC cell type (ON red, OFF blue, ON-OFF magenta, and Null grey), and the size of the dot indicates if the cell is core (big dot) or non-core (small dot) for each ensemble. ON ensembles are dominated by ON and ON-OFF cells, while the OFF ensembles are dominated by OFF and ON-OFF cells. **(C)** For each ensemble, the average PSTHs of their corresponding core-cells grouped by cell type shows no clear tuning preference. **(D)** The PSTH for each ensemble shows the stimulus preference of the ON and OFF ensembles. The different cell responses can be classified as transient and sustained within each ensemble, showing detailed stimulus tuning. **(E)** The colored matrix shows the RGCs in rows and the ensembles in the columns. Each column is colored in the entries corresponding to their core-cells, according to which ensemble they belong (A).

These different evoked responses led us to expect at least four types of ensembles: two ensembles related to transitions (one for the ON-OFF and one for the OFF-ON) and two ensembles related to the decaying-stable activity after the ON-OFF and OFF-ON transition, respectively. To test this hypothesis, we applied our ensemble detection algorithm (see Methods for details and parameters) on the spiking activity of 319 RGCs during 120 seconds of MEA recording.

We found ten ensembles comprising *∼* 68% of the spike patterns in the analyzed recording. Their activity was highly locked to the stimulus (Fig. 1A, D, middle and bottom panel). We found two transiently active ensembles (one for the ON-OFF and one for the OFF-ON transition), whose activity was only evoked by the stimulus transition, showing no activity either before the stimulus start (black arrow in Fig. 1A) or during the decaying or stable response. The other eight ensembles were active before stimulus presentation, but at a lower rate, and during the decaying or stable response. Notably, four of them (Ens. 7-10) are preferentially active in the OFF stimulus, and the other four during the ON stimulus (Ens. 2-5).

These results show that the detected retinal ensembles are preferentially tuned to the features of the stimulus, showing that without any stimulus-related information, our method can obtain ensembles whose activity is functionally coupled with the stimulus.

### Functional properties of RGC ensembles

We detect the ensembles and their *core-cells* i.e. cells whose correlation with a given ensemble is statistically significant(see Methods for details). On average, each core-cell participated in 2.7 *±* 1.3 ensembles, and only four RGCs were considered non-core, considering all the ensembles. Three cells participated in six ensembles, indicating that some cells may participate in both ensemble classes.

The two transiently active ensembles, namely Ens. 1 (ON) and Ens. 6 (OFF), have 257 and 222 core-cells, respectively, while the rest of the ensembles have, on average, 47.2 *±* 17.9 core-cells (Fig. 1E). This result is consistent with the increased population activity evoked by stimulus transition, where many cells are firing, and decaying responses where many cells become silent.

Using the RGC responses to the repeated ON-OFF stimulus (20 trials) and an automated RGC functional classification algorithm (see Methods for details), we obtained 44 ON, 23 OFF, 205 ON-OFF, and 47 Null (no significant preference) RGCs.

All the ensembles were composed of ON-OFF core-cells, but the ON ensembles were dominated by ON RGCs while OFF ensembles by OFF RGCs, as shown by the spatial distribution of the RGCs receptive field centers in Fig. 1B (see Methods for details). Null cells showed significant participation in some ensembles, despite their lack of preference according to the classification algorithm Then, in this simple setup, we found that the ensemble’s classification is consistent with their corresponding core-cells’ dominant classes.

The detection of core-cells allows us to investigate if the tuning preferences of ensembles are inherited from their core-cells or if the ensembles have a specific tuning preference as a functional unit. To evaluate this for each ensemble, we averaged the peri-stimulus time histogram (PSTH) of their corresponding core-cells grouped by RGC class Fig. 1C, obtaining one average PSTH per cell class.

Since ON-OFF cells are present in all ensembles, the core-cells of each ensemble, as a group, have a preference for both light transitions, despite the precise stimulus tuning of the ensembles (Fig. 1D).

The PSTH for each ensemble was also computed to provide the detailed tuning of each ensemble and compare the core-cells tuning preferences and the ensemble tuning preference.

We conclude that the ensemble tuning preference cannot be completely derived from their core-cells’ tuning preference, providing evidence in favor of neuronal ensembles as whole functional units.

### Method overview

Before a systematic evaluation of the performance of our method, we summarize its core elements using the example shown in Fig. 2, and refer the reader to the Methods section for further details.

**Fig 2.**
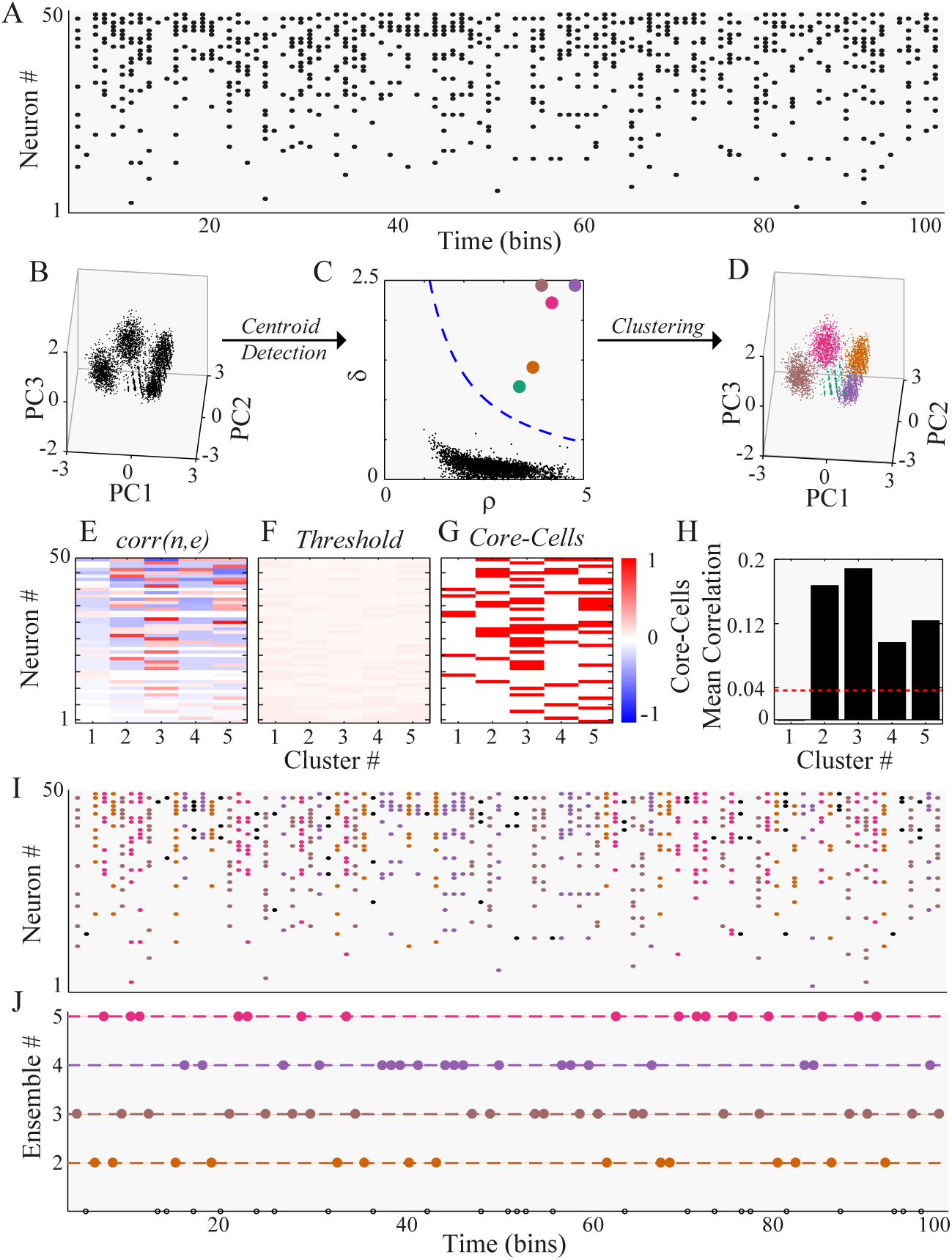
Methodological scheme. **(A)** Synthetic spike trains of 50 neurons during 100-time bins. Four ensembles were artificially generated (see Methods for details). **(B)** PCA is performed on all the spike patterns of more than three spikes. For visualization purposes, we plot each of them as a point in the 3D space spanned by the first three principal components (PC). The points tend to group in clusters. **(C)** The Euclidean distance between all the points on the PC coordinates is computed. The density of each data point *ρ* and its distance to the next data point with a higher density of *δ* is obtained. The points above the threshold (blue dashed line, see Methods for details) are considered cluster centroids. **(D)** All points are assigned to their closest centroid, building up the clusters of spiking patterns. **(E)** The correlation matrix between neurons and clusters, *corr*(*n, e*) is represented by the symmetric red-blue color map. **(F)** A threshold matrix is computed to define the significance of *corr*(*n, e*) values. **(G)** Core-cells are detected based on *corr*(*n, e*) values that exceed the threshold (red if core, white otherwise). **(H)** The inner-cluster mean correlation is compared against a threshold of non-core cell correlation, discarding any cluster below this threshold (cluster 1, in this case). This final filtering yields the set of ensembles. **(I)** Each spike pattern shown in **A** is colored according to the cluster to which they belong. Black patterns were discarded by the minimum number of spikes or the threshold rejection criteria. **(J)** The detected ensemble sequence is represented as an integer sequence, where the color corresponds to the spike pattern color in **I**. Note that the rejection criteria discarded cluster 1, so only ensembles 2-5 are shown, and black circles denote non-clustered spike patterns.

First, we discard any population pattern of the spike trains (Fig. 2A) with less than three spikes, and then perform a principal component analysis (PCA) using the population patterns as observations (Fig. 2B). Then, we computed the Euclidean distance between all the patterns projected on a small subset of the first principal components (from three to six), generating a distance matrix, from which the local density, *ρ*, and the distance to the next denser point, *δ*, is computed for each population pattern. Based on these two measures, and a power-law fit to the *ρ* vs. *δ* curve, we automatically detect the cluster centroids (Fig. 2C), and assign the rest of the patterns to their closest centroid, building up the clusters (Fig. 2D). Since the core of our algorithm is the density-based clustering procedure, we call it *Density-based*.

To find the core-cells, we computed *corr*(*n, e*), the correlation between the spike train of neuron *n* and the activation sequence of cluster *e* (Fig. 2E), and test its significance using a threshold associated with a null hypothesis obtained from shuffled versions of data (Fig. 2F). If *corr*(*n, e*) is above the threshold, neuron *n* is considered a core-cell of cluster *e* (Fig. 2G).

Finally, to obtain the ensembles, the pairwise correlation between all the core-cells of a given cluster are computed and averaged, representing the within-cluster average correlation.

To define a cluster be an ensemble, we compared the within-cluster average correlation to the average pairwise correlation of the whole network (Fig. 2H). If the within-cluster if significantly higher (based on a threshold), we considered an *ensemble* to be present; otherwise, the cluster is discarded due to the lack of internal correlation.

With this procedure, we were able to split the population patterns into an ensemble and non-ensemble patterns (Fig. 2I), and to obtain the activation sequence of different ensembles in time (Fig. 2J) with their corresponding core-cells.

### Detection of ensembles on synthetic data

To test our method, we designed an algorithm to generate synthetic data where neuronal ensembles’ activity can be parametrically controlled (see Methods for details). We assessed the detection performance of our method concerning known ground truth (GT). We illustrate our results with a simple example shown in Fig. 3, where different spike trains were generated using a network of *N* = 100 neurons and *T* = 5000 bins (the figure shows just 200 bins to improve the visualization). We created seven ensembles defining the temporal sequence of the ensembles, whose global temporal activity comprised 80% of the sample (Fig. 3E), and the core-cells, whose participation comprised from 20 to 40 neurons (Fig. 3I). Then, we randomly added or removed spikes to each neuron in order to satisfy a given firing probability for each neuron, *P* (*n*), which controls the spike trains density (Fig. 3B, C, D).

**Fig 3.**
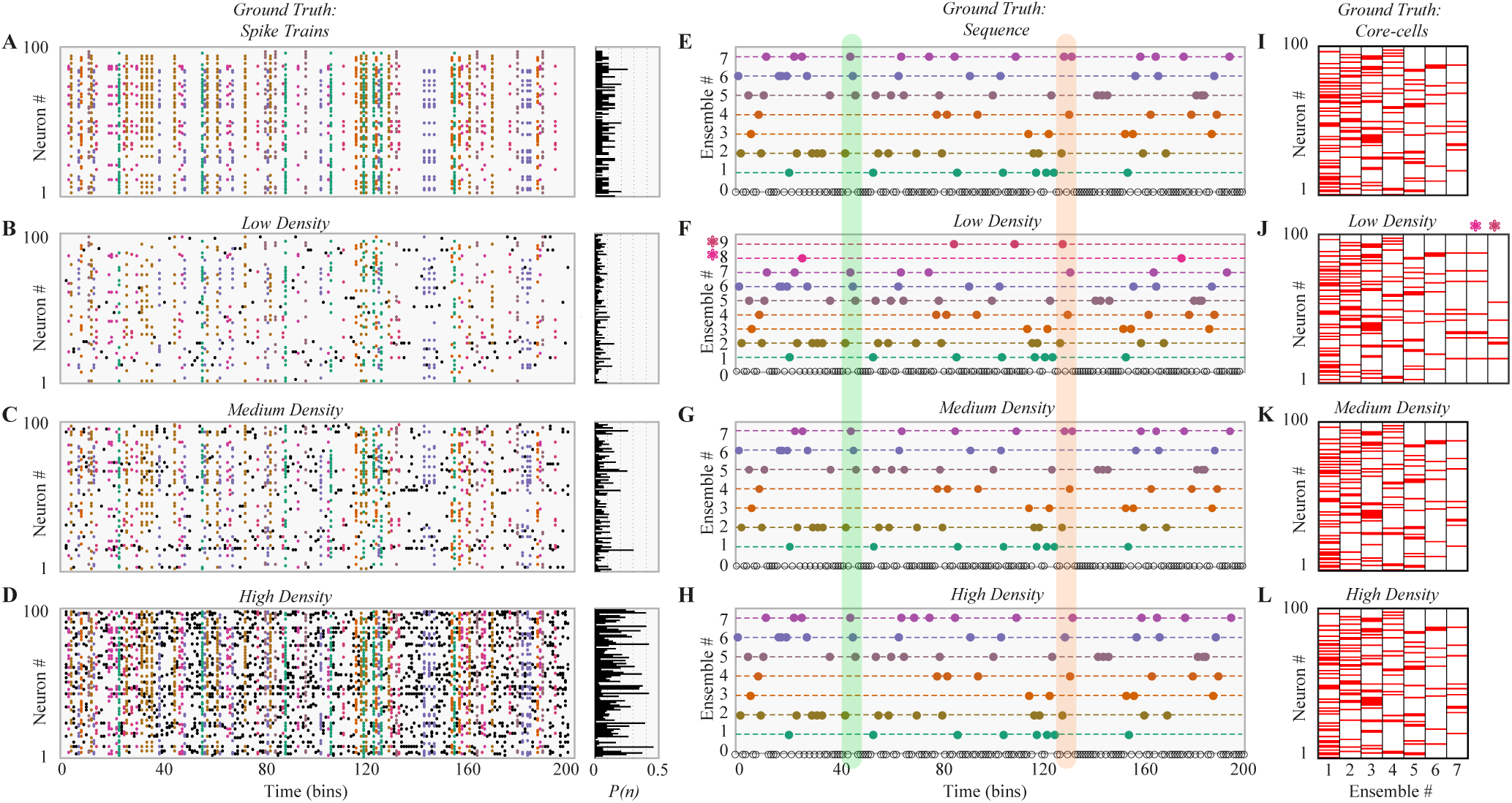
Robust ensemble detection on synthetic data under different density regimes. **(A)** The spatial (core-cells) and temporal (sequence) properties of seven ensembles are synthetically generated on a spike train of 100 neurons. The right panel shows the firing probability of each neuron at each time bin denoted by *P* (*n*). Each ensemble is colored differently. **(B-D)** Based on the ground truth (GT) spike trains, three spike trains with different densities were generated: low **B**, medium **C** and high density **D**. The firing probability of each neuron is shown at the right as in **A**. Spike patterns are colored as in **A**. Black dots are spike patterns that do not belong to any ensemble. **(E)** Temporal sequence of the GT ensembles. Color corresponds with **A**. **(F-H)** The detected ensemble sequence for the three spike trains with different densities, sorted to match the GT indexing and their colors. At low density, the method detects two extra ensembles, denoted by a colored asterisk. The green vertical region shows an example of ensemble that was correctly detected for the three spike trains. The red vertical region shows an example where the method correctly detects the GT pattern for Medium density, but partially fails in the other two cases. **(I)** GT core-cells are sorted in descending order according to the number of core-cells. Each column represents an ensemble, where red indicates a core-cell. **(J-L)** Detected core-cells for the three cases. Despite the two extra ensembles found in **J** (8 and 9, asterisks), core-cells are in good agreement with GT.

We found two ensembles more than expected (seven) for the low-density spike-trains (Fig. 3F, red asterisks), while for the medium and high-density ones we found the expected number of ensembles (Fig. 3G, H, respectively).

Then, we compared the ensemble sequence of the GT (Fig. 3E) and the detected ensemble sequence in each density scenario (Fig. 3F, G, H), finding almost perfect agreement between both, with the exception of a few false positives in the case of low and high density. Finally, we compared the detected core-cells with the GT (Fig. 3J, K, L), finding good agreement between both. Despite the over-detection in the low-density regime (red asterisk), the other ensembles were in good agreement with the GT core-cells.

With this example, we show that our method can detect the ensemble number, the temporal sequence, and the core-cells of neuronal ensembles in noisy spike trains with different densities. In the next section, we evaluate the detection of three features, i.e., ensemble number, temporal ensemble sequence, and core-cells, for a different sample and network sizes.

### Scaling performance and comparison with an alternative method

To systematically quantify and evaluate the performance of our method (Density-based), we generated synthetic data with different network and sample sizes and compared our algorithm to a current state-of-the-art ensemble detection method published by Carrillo *et al.* (SVD-based) [16]. Due to the latter method’s computational cost, we only compared the scaling with the sample size for both methods, while for the former, we also computed the scaling of performance with network size. Both methods were applied using default parameters provided by their respective computational codes.

First, we generated a synthetic spike train with fixed network size (*N* = 300), number of ensembles (*E* = 12), number of core-cells equal to 35, ensemble probability (*P*_*E*_ = 0.8), and medium density. Then, we varied the recording length from *T* = 500 to *T* = 10^4^, finding that both methods increased the computational time with the sample size, as expected, but the SVD-based method scaled exponentially, while the Density-based method is two orders of magnitude faster for *T* = 10^4^ (Fig. 4A).

**Fig 4.**
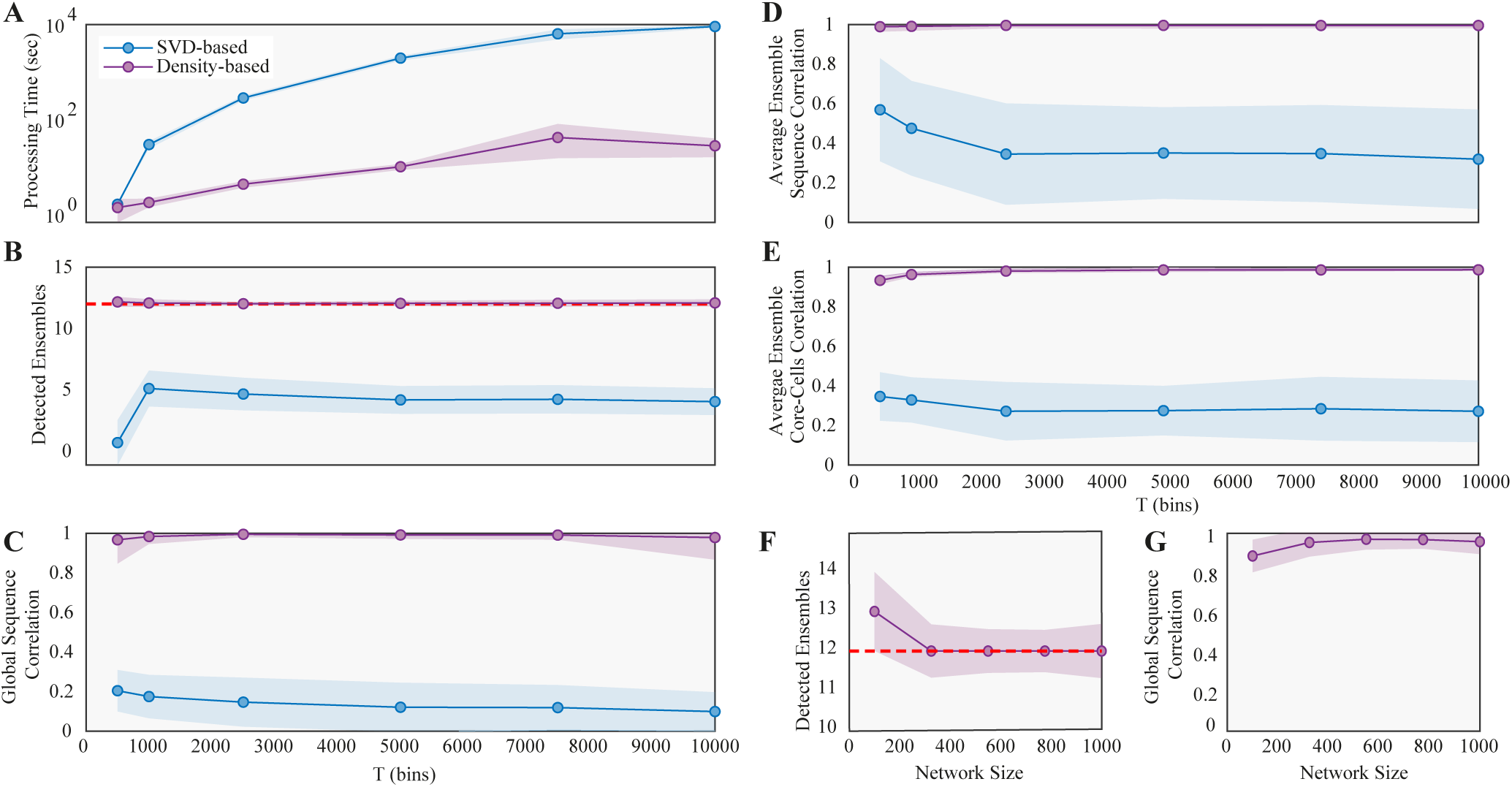
Performance scaling with sample size and network size. **(A)** Computational processing time in log scale of our method (purple) and Carrillo et al [16] (blue), as a function of sample size T (bins). Synthetic data generated using *N* = 300, medium density (see Methods for details) and 35 neurons per ensemble. Dots are averages, and shaded regions are *±* 1 standard deviation of 100 repetitions. **(B)** Number of detected ensembles. The dashed red line is the real Number of ensembles. **(C)** Correlation between the detected and true global ensemble sequence. **(D)** Average correlation between detected and true individual ensemble sequence, and **(E)** between detected and true ensemble core-cells. **(F)** Number of detected ensembles as a function of network size. The Number of core-cells per ensemble corresponds to the 35% of the network size. **(G)** Same as **C**, but as a function of network size. Due to computational cost, only our method was evaluated as a function of network size *T* = 5000.

Regarding the ensemble activity, our method accurately detects the number of ensembles for samples as small as *T* = 1000, while the SVD-based method converges to underestimation of the ensemble number (Fig. 4B). Once the detected ensembles were matched to their closest GT ensemble (see Methods for details), we measured the correlation between both sequences, finding that our method detects the GT with excellent performance over the whole range of sample sizes. The SVD-based method, in turn, systematically fails to detect the global sequence (Fig. 4C).

Furthermore, we measured the average correlation between the detected and GT individual sequences, and between the detected and GT core-cells, finding again that our method achieved excellent performance for both features even for small sample sizes (Fig. 4D, E, respectively).

Finally, to evaluate the performance for different network sizes, we fixed the sample size, while also keeping the other parameters fixed (number of ensembles, of core-cells and *P*_*E*_), and varied the network size from *N* = 50 to *N* = 1000, finding that our method slightly overestimated the ensemble number for small *N*, but yielded accurate results for the larger *N* (Fig. 4F, G).

We conclude that our method reliably performs on a wide range of sample and network sizes, giving a more scalable and accurate solution to the ensemble detection problem than the alternative SVD-based method.

### Reliable performance over a wide range of spike-train parameters

Here, we show the performance of our method when the number of ensembles and the number of core-cells vary independently.

We generated synthetic spike-trains with fixed network size (*N* = 300), sample size (*T* = 5000), ensemble probability (*P*_*E*_ = 0.8), and medium density while the number of ensembles and core-cells varied, as shown on Fig. 5. We found a wide combination of these parameters where the method detects the number of ensembles with a small relative error (Fig. 5A), accurately detects the global ensemble sequence (Fig. 5B), and the corresponding core-cells (Fig. 5C). Further explorations of other parameters and combinations of the same are considered work to be developed. To this end, we provide the computational codes and a GUI at https://github.com/brincolab/NeuralEnsembles. This GUI allows one to perform all the analyses presented here on multi-variate recordings of single events (e.g., spiking data, calcium events, arrival times in a sensor).

**Fig 5.**
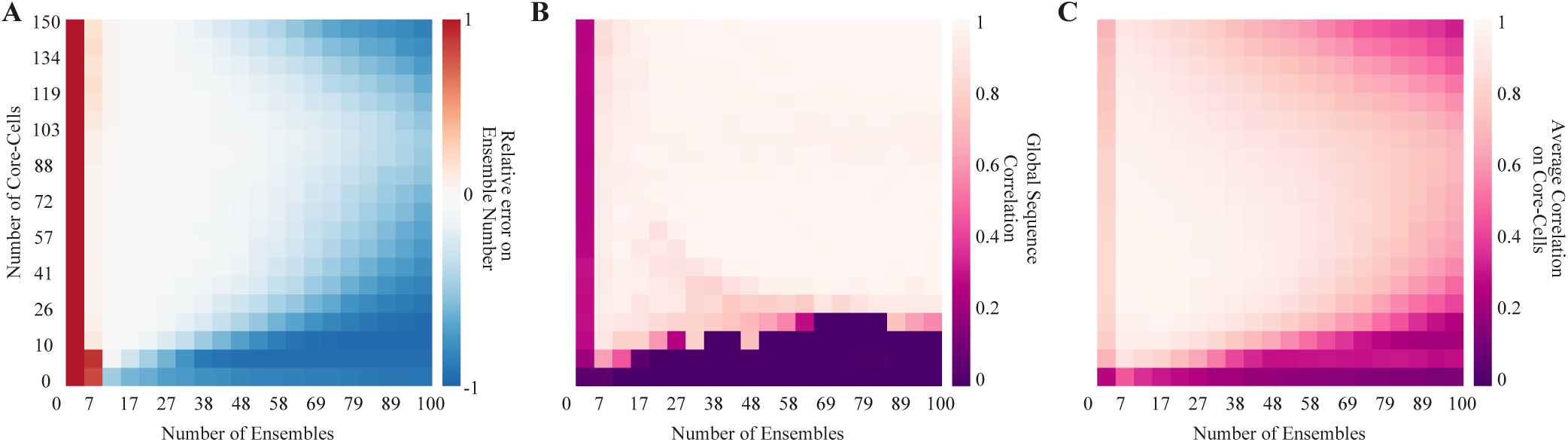
Detection performance over a wide range of spike-train parameters. **(A)** The relative error of detecting the ensemble number for spike-trains with different ensembles and the different number of core-cells. Synthetic data generated using medium density, *N* = 300, and *T* = 5000. Heat maps represent the average over 100 repetitions using the same spike-train parameters. Red/blue represents over/under detection respectively. **(B)** The correlation between the detected and the GT ensemble sequence. **(C)** Average correlation between the detected and the GT core-cells.

## Materials and methods

### Synthetic Spike Trains

We consider binary spike trains arranged in a matrix **S**_*N×T*_ where *N* corresponds to the number of neurons and *T* the number of time bins. The entries of the matrix denoted by **S**_*n,t*_ are equal to one of the *n*-th neuron is active on the *t*-th time bin, and zero otherwise. At each time bin *t*, there is a binary population pattern of active and silent neurons (**S**_1,*t*_, …, **S**_*N,t*_).

We generated a set of ensembles *E* characterized by their *core-cells* and an *activation sequence* of ensembles. Each ensemble was composed of a fixed number of core cells randomly drawn from the whole population of neurons, allowing the repetition of core-cells among ensembles. We generated for each ensemble, a column binary vector *c*_*e*_ of dimension *N*, with *c*_*e*_(*n*) = 1 if neuron *n* is a core-cell of ensemble *e* and 0 otherwise.

The activation sequence of each ensemble *e ∈ E* follows a time homogeneous Bernoulli process with parameter *P*_*E*_, denoted *B*(*P*_*E*_). We generate a row binary vector denoted by *a*_*e*_ of dimension *T* with *a*_*e*_(*t*) = 1 if ensemble *e* is active on time bin *t* and 0 otherwise. We allowed at most one active ensemble per bin. To implement this idea, we first drew the first ensemble’s activation times at random and then removed these times from the list of possible time points available for the second ensemble. We proceeded this way until reaching *P*_*E*_.

With the core-cells and the activation sequence of each ensemble defined, a spike train was generated by the product *c*_*e*_*a*_*e*_ (matrix) of dimension *N × T*.

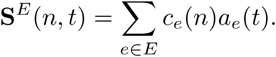

In order to preserve the probabilistic nature of spiking neurons, the firing rates of each neuron was drawn from a rectified Gaussian distribution (only positive values) with variable standard deviation (*s.d.*). The larger the *s.d.*, the higher the firing rates are present on the spike train. This parameter allowed for the control of the density of the spike train (the total number of spikes respect to the spike train duration).

We randomly added/removed spikes to/from each neuron’s spike train until matched the target firing rate *P* (*n*). In removing spikes, the spike pattern located on the time bins where a given ensemble was active, end up having less active neurons than defined; on the other hand, if we added spikes, the opposite effect occurred. We used three different spike train densities: low (*s.d.* = 0.05), medium (*s.d.* = 0.1) and high (*s.d.* = 0.2). For the low-density case, matching the target firing rates usually required the removal of spikes. This method corrupted the patterns related to ensemble activity by underrepresenting their core-cells. In the medium case, we had a balance between adding and removing spikes, and, in the last case, most neurons required the addition of spikes to match the desired firing rate, making neurons participate in ensembles they did not belong by construction.

In summary, we generated a spike train from the following procedure:

1. Generate *E* ensembles characterized by spike patterns built from their core-cells.
2. Fill a spike train with the patterns of active ensembles following a time-homogeneous Bernoulli process for each ensemble considering the proportion of *P*_*E*_ of the complete set of spike patterns. The remaining spike patterns are filled with the spike pattern consisting only of silent neurons.
3. Draw the firing rates of each neuron from a rectified positive Gaussian distribution.
4. Randomly add/remove spikes to/from each neuron’s spike train until it matches the target firing rate of each neuron.

This procedure yields a random spike train built from the ensemble activity.

## Ensemble Detection

### Feature Extraction using PCA

To detect neuronal ensembles from data, we used Principal Component Analysis (PCA) over a subset of population patterns. PCA extracts a set of orthogonal directions capturing the most significant variability observed on the spike patterns. We discarded the spike patterns with less than three spikes and projected the spike patterns on the first six principal components (PCs). Using four to seven principal components yielded no difference in the clustering procedure. However, the optimal number of PCs may vary depending on the data, so it is considered a free parameter in the GUI. We computed the Euclidean distance between all binary patterns projected on the first six principal components to guide close patterns’ clustering.

### Centroid detection

Two parameters characterized each data point (i.e., a population pattern): i) *ρ* the density and ii) *δ* the minimum distance to a point with higher density. The density of each pattern *i* was given by 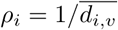, where 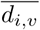 was the average distance from point *i* to its closest neighbors. Typically, we considered 2% of the closest neighbors. This choice can be tuned with a parameter we denoted *d*_*c*_. Spike patterns with relatively high values of *ρ* and *δ* were candidates of centroids. The rationale is that points with relatively high *ρ* and *δ* (respect to the rest of the points) have the highest number of points in their vicinity and are far from other points with high density. The procedure to find the centroids of the clusters follows [41], which is a modified version of the method developed in [42]. To automatically detect the cluster centroids, we fit a power-law to the *δ* vs. *ρ* curve, using the 99.9% upper confidence boundary as a threshold. We considered centroids to be all points falling outside this boundary. The rest of the points were assigned to their closest centroids. Thus, each binary pattern with more than three spikes was assigned to a cluster, obtaining, in this way, their corresponding activation times.

### Core-cell detection

Once the candidate clusters and centroids were identified, the Pearson correlation coefficient between the activation times of neuron *n* and cluster *e*, denoted by *corr*(*n, e*), was computed for each pair (*n, e*). This correlation is 1 if neuron *n* and cluster *e* were always active in the same time bins, and is -1 if they are never active at the same time bins. The intermediate values represented combinations of neurons and clusters with a tendency to be active either in the same or in different time bins.

To distinguish between a core and a non-core cell, we set a threshold obtained from a null hypothesis built from shuffled versions of the (*n, e*) pair. For each cluster, *e ∈ E*, 3 the number of times in which it was active is kept fixed, but the temporal sequence was randomly shuffled, obtaining a new temporal sequence of activation that we denoted *e*_*r*_.

We repeated this procedure 5000 times for each cluster, obtaining a distribution of *corr*(*n, e*_*r*_) for neuron *n* in cluster *e*, which represented the null hypothesis distribution associated with the correlation between a neuron and a cluster given by chance. A significance threshold was then defined as the 99.9-th percentile of this distribution; therefore, we set *p <* 0.001 as threshold *θ*_*e*_. Then, if *corr*(*n, e*) *> θ*_*e*_, we considered that *n* was a core-cell of cluster *e*, as their correlation was above the significance level.

### Ensemble selection criteria

Following [43], we used two criteria: i) minimum cluster size and ii) within-cluster average correlation. The first criterion considered as candidates for neuronal ensembles clusters whose number of core-cells was above three. The second criterion compared the average pairwise correlation between all the core-cells of the same cluster to the average pairwise correlation of the whole population plus a threshold. This way, we kept only clusters with a minimum size (in terms of core-cells) and with relatively (concerning the population) high within-cluster average correlation. If a cluster failed to pass any of these two criteria, we discarded it from the analysis. Otherwise, we considered the cluster as an ensemble.

### Matching detected ensembles with ground truth ensembles

To evaluate our method with synthetic data, we matched the GT ensembles with the detected ones. This was implemented by computing the correlation between the GT’s activation sequence and the detected ensembles. Thus, we looked for the detected ensemble that maximized the correlation with the GT ensembles.

## Discussion

We introduced a scalable and computationally efficient method to detect neuronal ensembles from spiking data. Using simple clustering techniques [42], and suitable statistical tests, we were able to develop a simple, fast and accurate method. On the one hand, our example of mouse retinal ganglion cells provides evidence for an expected causal relationship between stimuli and RGC ensembles. On the other hand, the simulated data examples show that our method provides accurate results for a wide range of neural activity scenarios, outperforming existing tools for ensemble detection in terms of accuracy and computation time.

Our method detects neural ensembles considering three general properties of them: transient activation (spontaneous and/or stimuli-evoked), the same neurons may participate in different ensembles, and the correlation between the core-cells (within-ensemble correlation) of a given ensemble should be high, compared to the rest of the population. Other methods for ensemble detection fulfill other criteria, e.g., finding communities between spiking neurons along time [26, 44], or using event-related activity [43]. Despite the usefulness of these methods in their context, our analysis is more general. It relies on grouping the population spiking patterns with no event-related information, allowing us to segment a spiking network’s temporal activity under both spontaneous and stimuli-evoked conditions.

Regarding the SVD-based method [9], we acknowledge the insights that its application has provided to the study of neural networks. However, it has limitations in terms of computational cost (for relatively small sample sizes, Fig. 4) and parameter tuning. Further, the SVD-based method extracts the temporal sequence of ensembles from the spectral decomposition of the similarity matrix between population patterns, aiming to detect groups of linearly independent patterns. Instead, we use a subset of the first principal components to embed the population patterns and cluster them into that space, with no need to find linear independence between the clusters. Finally, the SVD-based method has no explicit implementation for the evaluation of the within-ensemble correlation, which, in our case, is a critical step to distinguish between a noisy cluster and an ensemble cluster.

There is room for further improvement of our method. For example, non-linear spike correlations may produce spurious results in ensemble detection methods that depend on PCA. Alternative approaches can be adapted to other measures of spike train similarity besides linear correlations [45]. In our example of RGCs, while we use the standard bin size for mammalian RGCs, the bin size used to create the spike trains may influence the detected ensembles, and thus different bin sizes can yield different results.

Finally, we provided novel evidence in favor of the existence of retinal ensembles that are functionally coupled to the stimuli. However, our purpose was to exemplify our method on a biological spiking network rather than explaining the possible mechanisms involved in the activity of retinal ensembles or their functional implications. Indeed, we consider the study of retinal ensembles as an exciting new research avenue that needs to be developed in the way to understand vision, sensory systems, and more generally, the nervous system. In line with this perspective, we delivered a method that is general enough to be applied to any multi-variate binary data set. Furthermore, it is intuitive and can generate results that are easy to visualize, which should favor their comprehension and general use by the scientific community. Matlab codes and a GUI implementing our new method accompany this article.

## Supplementary Material

### Code

The GUI and the codes used to validate the method using synthetic data can be found at https://github.com/brincolab/NeuralEnsembles.

### Ethics Statement

Animal manipulation and breeding and corresponding experiments were approved by the bioethics committee of the Universidad de Valparaiso, in accordance with the bioethics regulation of the National Agency for Research (ANID, Ex-CONICYT) and international protocols.

### Animals and RGC Recordings

The experimental mice were maintained in the animal facility of the Universidad de Valparáıso, at 20–25°C on a 12-h light-dark cycle, with access to food and water *ad-libitum.* These recordings were performed for other experimental purposes, and for the present work only one recording was used. The corresponding methods of MEA recording have previously been described [46]. In brief, animals were euthanized under deep isofluorane or halothane anesthesia and both eyes were extracted. Then, one of the extracted retinas was diced into quarters while the other was stored in oxygenated (*O*_2_ 95% *CO*_2_ 5%) AMES medium at 32°C in the dark for further experiments. The same AMES media was used for continuous perfusion during extracellular recordings. For MEA recordings (MEA USB-256, 20kHz sample, Multichannel Systems GmbH, Germany), one piece of retina was mounted onto a dialysis membrane then placed into a ring device mounted in a traveling (up/down) cylinder, which was moved to contact the electrode surface of the MEA recording array. Data were processed off-line using the Spiking-Circus spike sorting algorithm [41] with default parameters.

### Visual Stimuli

Visual stimuli were generated by a custom software created with PsychoToolbox (Matlab) on a MiniMac Apple computer and projected onto the retina with an LED projector (PLED-W500, Viewsonic, USA) equipped with an electronic shutter (Vincent Associates, Rochester, USA) and connected to an inverted microscope (Lens 4x, Eclipse TE2000, NIKON, Japan). The image was conformed by 380 x 380 pixels, each covering 5*µm*^2^. To estimate the RGC receptive fields, a checkerboard stimulus (visual white noise) with a block size of 50*µm* was presented at a rate of 60 Hz for 20 mins, with each block independently taking 0 or 255 (max value) in the pixel value scale. To classify the RGC, a green ON-OFF light stimulus was presented, where each part lasted three seconds, repeated 21 times. For the classification analysis, the first trial was discarded.

### Automated RGC classification

RGCs were automatically classified as ON, OFF, ON-OFF, and Null depending on their preference to the light stimulus, using the statistical approach presented in Ref [47]. We computed the peri-stimulus time histogram (PSTH) for each RGC and compared the maximum activity in the ON and OFF part of the stimulus. Then, we set a threshold based on the average of the PSTH plus 2 *s.d.*, and if only the maximum in the ON (OFF) part was above this threshold, we considered this RGC as ON (OFF); if both maxima were above the threshold, we considered the RGC as ON-OFF. Otherwise, the RGC was classified as Null due to its lack of preference for the stimulus.

### RGC receptive fields estimation

The spike-triggered average (STA) of each RGC, defined as the average stimulus preceding a spike, was computed by the reverse correlation method using the checkerboard stimulus (see Methods subsection 2) aggregating the 18 frames previous to any emitted spike in a matrix as in Ref [48]. This STA matrix was decomposed using SVD, which estimates the temporal and spatial components of the receptive field, where the former represented the average stimulus fluctuation previous to a spike. In contrast, the latter represented the preferred location of the RGC in the stimuli space. Then, an ellipse was fitted to the spatial component to estimate the RGC receptive field.

## Acknowledgments

The authors thank Fernando Rosas for insightful discussions and valuable suggestions. Funding by CONICYT scholarship CONICYT-PFCHA/Doctorado Nacional/2018-21180428 (RH) and CONICYT scholarship CONICYT-PFCHA/Maǵıster Nacional/2020-22200156 (AM). Fondecyt Iniciacíon 2018 Proyecto 11181072 (RC). AFOSR Grant FA9550-19-1-0002 (MJE and AGP). ICM-ANID P09-022-F, CINV (AGP).

## References

1. Hebb DO. The Organization of Behaviour. Organization. 1949; p. 62. doi:citeulike-article-id:1282862.

2. Russo E, Durstewitz D. Cell assemblies at multiple time scales with arbitrary lag distributions. eLife. 2017;6(1):e19428. doi:10.7554/eLife.19428.

3. Varela F, Lachaux JP, Rodriguez E, Martinerie J. The brainweb: Phase synchronization and large-scale integration. Nature Reviews Neuroscience. 2001;2(4):229–239. doi:10.1038/35067550.

4. Uhlhaas P, Pipa G, Lima B, Melloni L, Neuenschwander S, Nikolíc D, et al. Neural synchrony in cortical networks: history, concept and current status. Frontiers in Integrative Neuroscience. 2009;3:17. doi:10.3389/neuro.07.017.2009.

5. Stringer C, Pachitariu M, Steinmetz N, Reddy CB, Carandini M, Harris KD. Spontaneous behaviors drive multidimensional, brainwide activity. Science. 2019;364(6437). doi:10.1126/science.aav7893.

6. Yuste R. From the neuron doctrine to neural networks; 2015.

7. Ringach DL. Spontaneous and driven cortical activity: implications for computation; 2009.

8. Carrillo-Reid L, Yuste R. Playing the piano with the cortex: role of neuronal ensembles and pattern completion in perception and behavior; 2020.

9. Carrillo-Reid L, Lopez-Huerta VG, Garcia-Munoz M, Theiss S, Arbuthnott GW. Cell Assembly Signatures Defined by Short-Term Synaptic Plasticity in Cortical Networks. International journal of neural systems. 2015;25(7):1550026. doi:10.1142/S0129065715500264.

10. Buzsáki G. Large-scale recording of neuronal ensembles; 2004.

11. Einevoll GT, Franke F, Hagen E, Pouzat C, Harris KD. Towards reliable spike-train recordings from thousands of neurons with multielectrodes. Current opinion in neurobiology. 2012;22(1):11–7. doi:10.1016/j.conb.2011.10.001.

12. Watanabe K, Haga T, Tatsuno M, Euston DR, Fukai T. Unsupervised detection of cell-assembly sequences by similarity-based clustering. Frontiers in Neuroinformatics. 2019;doi:10.3389/fninf.2019.00039.

13. Harris KD. Neural signatures of cell assembly organization. Nature reviews Neuroscience. 2005;6(5):399–407. doi:10.1038/nrn1669.

14. Carrillo-Reid L, Yang W, Kang Miller Je, Peterka DS, Yuste R. Imaging and Optically Manipulating Neuronal Ensembles. Annual Review of Biophysics. 2017;doi:10.1146/annurev-biophys-070816-033647.

15. See JZ, Atencio CA, Sohal VS, Schreiner CE. Coordinated neuronal ensembles in primary auditory cortical columns. eLife. 2018;doi:10.7554/eLife.35587.

16. Carrillo-Reid L, Miller JEK, Hamm JP, Jackson J, Yuste R. Endogenous Sequential Cortical Activity Evoked by Visual Stimuli. Journal of neuroscience. 2015;35(23):8813–28. doi:10.1523/JNEUROSCI.5214-14.2015.

17. Wenzel M, Hamm JP, Peterka DS, Yuste R. Acute Focal Seizures Start As Local Synchronizations of Neuronal Ensembles. The Journal of neuroscience: the official journal of the Society for Neuroscience. 2019;doi:10.1523/JNEUROSCI.3176-18.2019.

18. Hamm JP, Peterka DS, Gogos JA, Yuste R. Altered Cortical Ensembles in Mouse Models of Schizophrenia. Neuron. 2017;doi:10.1016/j.neuron.2017.03.019.

19. Wenzel M, Han S, Smith EH, Hoel E, Greger B, House PA, et al. Reduced Repertoire of Cortical Microstates and Neuronal Ensembles in Medically Induced Loss of Consciousness. Cell Systems. 2019;8(5):467 – 474.e4. doi:https://doi.org/10.1016/j.cels.2019.03.007.

20. Fang WQ, Yuste R. Overproduction of Neurons Is Correlated with Enhanced Cortical Ensembles and Increased Perceptual Discrimination. Cell Reports. 2017;doi:10.1016/j.celrep.2017.09.040.

21. Buzsáki G. Neural Syntax: Cell Assemblies, Synapsembles, and Readers. Neuron. 2010;68(3):362–385. doi:10.1016/j.neuron.2010.09.023.

22. Eichenbaum H. Barlow versus Hebb: When is it time to abandon the notion of feature detectors and adopt the cell assembly as the unit of cognition?; 2018.

23. Nicolelis MAL, Baccala LA, Lin RCS, Chapin JK. Sensorimotor encoding by synchronous neural ensemble activity at multiple levels of the somatosensory system. Science. 1995;doi:10.1126/science.7761855.

24. Benchenane K, Peyrache A, Khamassi M, Tierney PL, Gioanni Y, Battaglia FP, et al. Coherent Theta Oscillations and Reorganization of Spike Timing in the Hippocampal-Prefrontal Network upon Learning. Neuron. 2010;doi:10.1016/j.neuron.2010.05.013.

25. Peyrache A, Benchenane K, Khamassi M, Wiener SI, Battaglia FP. Principal component analysis of ensemble recordings reveals cell assemblies at high temporal resolution. Journal of Computational Neuroscience. 2010;doi:10.1007/s10827-009-0154-6.

26. Lopes-dos Santos V, Conde-Ocazionez S, Nicolelis MAL, Ribeiro ST, Tort ABL. Neuronal assembly detection and cell membership specification by principal component analysis. PLoS ONE. 2011;6(6). doi:10.1371/journal.pone.0020996.

27. Singh A, Humphries MD. Finding communities in sparse networks. Scientific Reports. 2015;5:1–7. doi:10.1038/srep08828.

28. Bruno AM, Frost WN, Humphries MD. Modular deconstruction reveals the dynamical and physical building blocks of a locomotion motor program. Neuron. 2015;86(1):304–318. doi:10.1016/j.neuron.2015.03.005.

29. Torre E, Picado-Muiño D, Denker M, Borgelt C, Grün S. Statistical evaluation of synchronous spike patterns extracted by frequent item set mining. Frontiers in computational neuroscience. 2013;7:132. doi:10.3389/fncom.2013.00132.

30. Torre E, Canova C, Denker M, Gerstein G, Helias M, Grün S. ASSET: Analysis of Sequences of Synchronous Events in Massively Parallel Spike Trains. PLoS Computational Biology. 2016;doi:10.1371/journal.pcbi.1004939.

31. Yegenoglu A, Quaglio P, Torre E, rün S, Endres D. Exploring the usefulness of formal concept analysis for robust detection of spatio-temporal spike patterns in massively parallel spike trains. In: Lecture Notes in Computer Science (including subseries Lecture Notes in Artificial Intelligence and Lecture Notes in Bioinformatics); 2016.

32. Quaglio P, Yegenoglu A, Torre E, Endres DM, Grün S. Detection and Evaluation of Spatio-Temporal Spike Patterns in Massively Parallel Spike Train Data with SPADE. Frontiers in Computational Neuroscience. 2017;11(May):1–17. doi:10.3389/fncom.2017.00041.

33. Onken A, Liu JK, Karunasekara PPCR, Delis I, Gollisch T, Panzeri S. Using Matrix and Tensor Factorizations for the Single-Trial Analysis of Population Spike Trains. PLoS Computational Biology. 2016;doi:10.1371/journal.pcbi.1005189.

34. Euler T, Haverkamp S, Schubert T, Baden T. Retinal bipolar cells: elementary building blocks of vision. Nature reviews Neuroscience. 2014;15(8):507–19. doi:10.1038/nrn3783.

35. Shekhar K, Lapan SW, Whitney IE, Tran NM, Macosko EZ, Kowalczyk M, et al. Comprehensive Classification of Retinal Bipolar Neurons by Single-Cell Transcriptomics. Cell. 2016;166(5):1308 – 1323.e30. doi:https://doi.org/10.1016/j.cell.2016.07.054.

36. Vlasits AL, Euler T, Franke K. Function first: classifying cell types and circuits of the retina. Current Opinion in Neurobiology. 2019;56:8 – 15. doi:https://doi.org/10.1016/j.conb.2018.10.011.

37. Masland RH. The Neuronal Organization of the Retina. Neuron. 2012;76(2):266 – 280. doi:https://doi.org/10.1016/j.neuron.2012.10.002.

38. Gollisch T, Meister M. Eye Smarter than Scientists Believed: Neural Computations in Circuits of the Retina. Neuron. 2010;65(2):150 – 164. doi:https://doi.org/10.1016/j.neuron.2009.12.009.

39. Real E, Asari H, Gollisch T, Meister M. Neural Circuit Inference from Function to Structure. Current Biology. 2017;27(2):189 – 198. doi:https://doi.org/10.1016/j.cub.2016.11.040.

40. Tikidji-Hamburyan A, Reinhard K, Seitter H, Hovhannisyan A, Procyk C, Allen A, et al. Retinal output changes qualitatively with every change in ambient illuminance. Nature Neuroscience. 2014;doi:10.1038/nn.3891.

41. Yger P, Spampinato GLB, Esposito E, Lefebvre B, Deny S, Gardella C, et al. A spike sorting toolbox for up to thousands of electrodes validated with ground truth recordings in vitro and in vivo. eLife. 2018;7:1–23. doi:10.7554/eLife.34518.

42. Rodriguez A, Laio A. Clustering by fast search and find of density peaks. Science. 2014;344(6191):1492–1496. doi:10.1126/science.1242072.

43. Montijn JS, Olcese U, Pennartz CMA. Visual Stimulus Detection Correlates with the Consistency of Temporal Sequences within Stereotyped Events of V1 Neuronal Population Activity. The Journal of Neuroscience. 2016;36(33):8624–8640. doi:10.1523/JNEUROSCI.0853-16.2016.

44. Humphries MD. Dynamical networks: finding, measuring, and tracking neural population activity using network theory. 2017;doi:10.1101/115485.

45. Humphries MD. Spike-train communities: Finding groups of similar spike trains. Journal of Neuroscience. 2011;doi:10.1523/JNEUROSCI.2853-10.2011.

46. Palacios-Muñoz A, Escobar MJ, Vielma A, Araya J, Astudillo A, Valdivia G, et al. Role of connexin channels in the retinal light response of a diurnal rodent. Frontiers in Cellular Neuroscience. 2014;8(August):1–13. doi:10.3389/fncel.2014.00249.

47. Carcieri S, Jacobs A, Nirenberg S. Classification of retinal ganglion cells: A statistical approach. Journal of neurophysiology. 2003;90:1704–13. doi:10.1152/jn.00127.2003.

48. Chichilnisky EJ. A simple white noise analysis of neuronal light responses. Network-computation in Neural Systems. 2001;12:199–213.

